# Life-history traits inform on population trends when assessing the conservation status of a declining tiger shark population

**DOI:** 10.1101/395509

**Authors:** Christopher J. Brown, George Roff

**Affiliations:** Australian Rivers Institute, Griffith University, 170 Kessels Road, Nathan, Queensland, 4111, Australia; School of Biological Sciences, University of Queensland, QLD 4072, Australia

**Keywords:** Bayesian model, informed prior, migratory species, IUCN red list, megafauna

## Abstract

The assessment of the conservation status of wide ranging species depends on estimates of the magnitude of their population trends. The accuracy of trend estimates will depend on where and how many locations within a species’ range are sampled. We ask how the spatial extent of sampling interacts with non-linear patterns in long-term trends to affect estimates of decline in standardised catch of tiger sharks (*Galeocerdo cuvier*) on the east coast of Australia. We apply a Bayesian trend model that uses prior information on life-history traits to estimate trends where we use data from all regions versus spatial subsets of the data. As more regions were included in the model the trend estimates converged towards an overall decline of 71% over three generations. Trends estimated from data only from northern regions or southern regions underestimated and overestimated the regional decline, respectively. When a subset of regions was modelled, rather than the full data-set, the prior informed by life-history traits performed well, as did a weakly informed prior that allowed for high variation. The rate of decline in tiger sharks is consistent with a listing East Coast Australia tiger sharks as endangered under local legislation. Monitoring programs that aim to estimate population trends should attempt to cover the extremes and mid-points of a population’s range. Life-history information can be used to inform priors for population variation and may give more accurate estimates of trends that can be justified in debates about the status of threatened species, particularly when sampling is limited.

## 1. Introduction

Determining the status of species threatened by human activities is important for informing on their management, including the investment of conservation funds and decisions about legal actions (Rodrigues et al. 2006). The status of a species is often defined on the basis of trends in population size (Conservation of Nature Species Survival Commission 2001). For instance, some Great Hammerhead Shark populations (*Sphyrna mokarran*) were listed as critically endangered on the International Union for the Conservation of Nature (IUCN) Red List, partly on the basis of an >80% population decline over the past three generations (Camhi et al. 2009). The performance of management actions aimed at averting decline should also be assessed by monitoring population trends (e.g. Ward-Paige et al. 2012). Monitoring data will be most useful when it covers sufficient spatial and temporal scales to estimate trends accurately.

Accurate estimates of the magnitude of a population change may be confounded by short-term and localised variation in abundance, or masked by measurement errors (Gaston & McArdle 1994). Local abundance measurements of mobile species may be biased by migration into and out of the sampling area (Forney 2000). Our ability to observe a species may also vary place to place because of environmental variation (e.g. Sköld & Knape 2018). This issue is likely to be worsened where sampling of abundances is limited to few locations within a species’ broader range, because fewer sites are more likely to exhibit random variations that do not reflect the population’s true trend (Forney 2000). The precision of population trend estimates will therefore depend on the spatial extent of sampling.

Appropriately formulated statistical models can separate short-term noise from important trends. Linear and log-linear models have been popular approaches to trend analysis (e.g. Baum & Blanchard 2010; Keith et al. 2015; Knape 2016). Linear models provide a simple phenomological model of trends, but may miss non-linear changes and be influenced by short-term temporal ‘outliers’ (Fewster et al. 2000; Knape 2016). Non-linear statistical models, like smoothing splines, can smooth over short-term deviations in abundance to capture longer-term non-linear trends (Fewster et al. 2000; Forney 2000; Knape 2016). A choice must then be made in the modelling about the degree of smoothing. This choice is usually made based on sample size (Fewster et al. 2000) or the level of smoothing is fitted empirically with a method like generalized cross validation (e.g. Knape 2016). However, empirical approaches to smoothing can lead to over-fitting and biased inferences on trends (Knape 2016).

A model’s fit to time-series data can also be controlled by using process models that explicitly account for species life-history traits (Kindsvater et al. 2018; Sköld & Knape 2018). Process models can be effective at discerning short-term noise from longer term trends driven by population dynamics (e.g. Wilson et al. 2011; Sköld & Knape 2018), but accurate estimation of population parameters can be difficult if the population trend exhibits a ‘one-way-trip’ (Szuwalski & Thorson 2017). ‘One-way-trips’ will be common in species data that are being analysed for extinction risk. What we need is an approach that takes strength from both the phenomological and process based approaches to obtain accurate estimates of population trends in the face of monitoring data that is limited in geographic and temporal extent.

Here we apply Bayesian hierarchical models to fit trends to population declines. We use a species’ life-history traits as Bayesian prior information to control the level of smoothing in the fitted trend line. The approach is thus a hybrid that blends phenomological description of trends with ecological processes that inform the smoothing. We first use simulations to explore the accuracy of fitted trends for species with a range of population growth rates, then as a case-study we modelled multi-decadal trends in a declining tiger shark population from the east coast of Australia (Holmes et al. 2012; Roff et al. 2018). We aimed to determine (A) how the choice of prior influences the models’ ability to detect a long-term trend in relative abundance; (B) how sampling fewer regions affects estimation of the long-term trend; (C) whether the appropriate choice of a prior can give a more accurate estimate of the large-scale trend when sampling is constrained to fewer regions; and (D) obtain statistically robust estimates of tiger shark population decline relative to the IUCN red list criteria.

## 2. Methods

### 2.1 Case-study

We analysed temporal variation in shark catch per unit effort in the Queensland Shark Control Program (QSCP, Kidston et al. 1992). The QSCP was instigated in 1962 at several sites around south-east Queensland, and has been expanded to 11 regions across 1760km of coastline, ranging from the tropics to sub-tropical areas (Figure 1). The QSCP uses a series of baited drumlines and mesh nets to capture sharks. The nets and drumlines are checked by contractors 15-20 days of each month, who also record the length and taxonomic identity of captured sharks. Effort data in the form of total number of nets and drumlines were reconstructed using historical records from contractor’s logbooks between 1962-2017. Estimates of effort were adjusted to reflect changes in protocols over time, including swapping of gear between beaches to avoid bycatch of whales (Roff et al. 2018). Where catch records were unclear or uncertainty existed regarding number of drumlines or nets, beaches were excluded from the analysis (Roff et al. 2018). Since the early 1990’s, drumlines and net types have been standardised across the program (Sumpton et al. 2010). A previous analysis of this same data-set found consistent declines across Queensland for the four major groups of sharks caught in the QSCP, including a 75% decline in tiger sharks over 1962-2017 (Roff et al. 2018).

**Figure 1.**
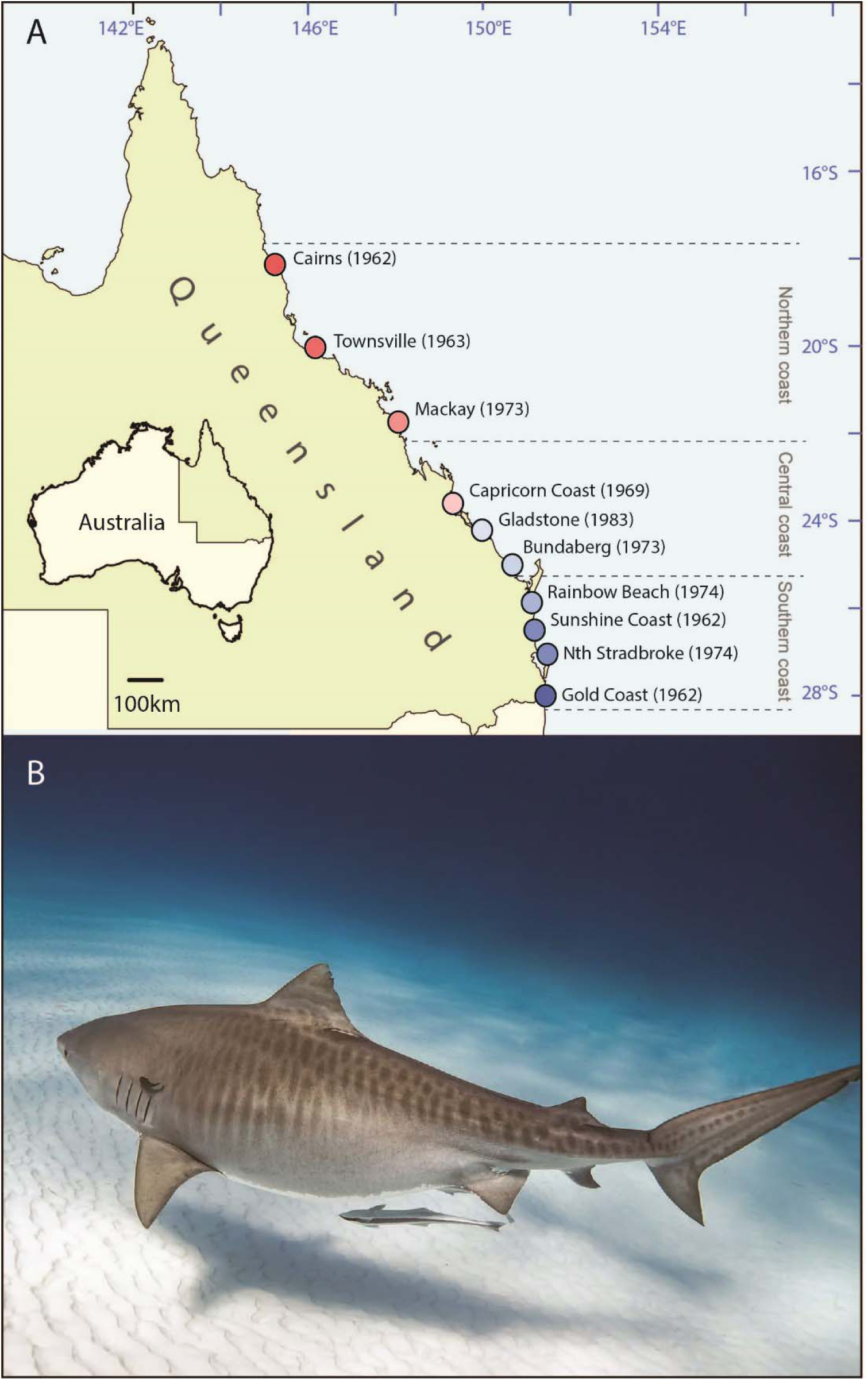
**A**) Map of the study region, showing major regions, first year of shark program sizes and scaling of sites from north to south, B) mature female *Galeocerdo cuvier* (∼3m length) showing characteristic vertical striping pattern. Photo credit: Juan Oliphant

Here, we explore the impact of model choice and the spatial extent of data on estimates of tiger shark declines. We focus on tiger sharks because: (1) they are reliably identified due to their distinctive body stripes (Fig 1b) (Holmes et al. 2012; Roff et al. 2018); (2) there are extensive catch records for this species, providing sufficient data for us to analyse the effect of using data from a sub-set of regions; (3) they may be of conservation concern because of the previously identified decline over 1962 and 2017 (Roff et al. 2018) and (4) the declines were non-linear, making tiger sharks a useful test-case for exploring the impact of non-linear trends on estimates of decline.

For inclusion in the IUCN red list, populations must exhibit observed, estimated, inferred or suspected population size reductions over three generations (Conservation of Nature Species Survival Commission 2001). Many regional agencies also apply the IUCN red listing criteria to determine local listing of species, including the state of Queensland and Australia (Committee 2019; Environment & Heritage Protection 2019). Analysis of sharks caught in the QSCP indicate an estimated age at 50% of maturity (A50) for female sharks of 10-13 years (Holmes et al. 2015). Based on this criteria, we analysed the QSCP dataset of 11 regions for trends in tiger shark catches between two time periods: three generations, which was 1984 to 2017 and the longer term trend over 1970 to 2017. 1970 was the earliest year shared by all sites used in the regional subsets.

### 2.2 Model

We used Bayesian hierarchical models to fit non-linear trend lines to tiger shark CPUE data. The complete model of the count of shark catch for each region, gear type and year (*y*_*i,g,t*_) was as follows(eqn 1):

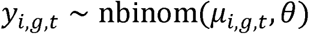

Where *θ* was the scale parameter of the negative binomial distribution and *µ*_*i,g,t*_ was the expected abundance. The expectation was specified (eqn 2):

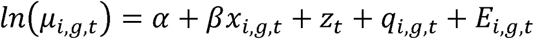

Where *α* was a global intercept, *β* was the additive effect of mesh nets, *x*_*i, g,t*_ was an indicator variable for nets (=1)or drumlines (=0) included to help control for differences in catches by different gear types (Holmes et al. 2012).

The east coast trend was estimated with *z*_*t*_, a latent first order random walk with mean and variance (Rue & Held 2005) (eqn 3):

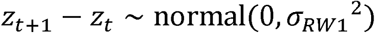

The random walk allows for non-linear trends in abundance with time, but will shrink toward a flat line for small values of *σ*_*RW*1,_ its standard deviation.

Region and gear specific devaitions from the east coast trend were estimated with an auto-regressivet erm (Sørbyea and Rue 2016) (eqn 4):

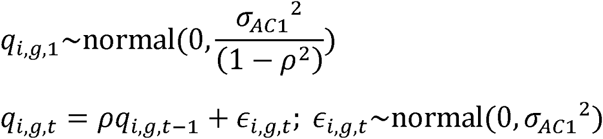

Where *ρ* is a correlation coefficient. When *ρ* = 0 the model reduces to a simple random effect on year/region/gear combinations. Together the random walk and auto-regressive components decompose variation among an east coast wide trend and region specific trends.

The term *E*_*i,g,t*_ was an offset term that had the logged number of nets or drumlines operating in each region and each year. The offset term means that we can re-arrange eqn 2 to present results in terms of CPUE.

To perform Bayesian computation we used the programming package INLA (Integrated Nested Laplace Approximation) (Rue et al. 2009, 2017) as implemented in the R programming environment (INLA version 17.06.20; R version 3.4.4; Martins et al. 2013; R Core Team 2018). Further details of INLA settings are explained in the Supplementary Material and code will be made available on Github upon publication.

### 2.3 Prior choice for the models

In these Bayesian models, the prior information is updated based on the data to estimate a posterior distribution for decline. For the parameters *α, β* we used broad normal priors. For the hyper-parameters *ρ* and *σ*_*AC1*_ we used the weakly informative priors that are the default in the INLA software (Supplementary Material). Using weakly informative priors means computations are more efficient and avoids overfitting of the data by the random effects (Simpson et al. 2017). We conducted a simulation study to choose a prior for *θ* (Supplementary Material).

We varied the priors for *σ*_*RW1*_ in simulations to address aims A and C. In random walk models, the prior for the standard deviation of the random walk interacts with real trends in the data to affect the level of smoothing in the trend line. A prior that has greater density closer to a smaller standard deviation will shrink the trend toward a constant line (Simpson et al. 2017). We used life-history information on the annual potential for variation in tiger shark abundance to inform the prior for the random walk. The standard deviation of a random walk has a direct interpretation in terms of population growth: For the exponential population model 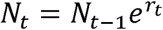, where *r*_*t*_ is an annually varying instantaneous growth rate and is sampled from a normal distribution, the standard deviation of a (logged) random walk will equal the standard deviation of r (*σ*_*RW1*_ = *σ*_*r*_). In applying this approach to CPUE data, we assumed that CPUE was proportional to population abundance.

One potential pitfall of the above approach is that normally distributed values of *r*_*t*_ imply the same probability of increases as decreases, which may result in the model missing large population declines. To overcome this problem we used a symmetrical prior that allows for large sustained changes in population size. The penalized complexity prior had a high density near a standard deviation of zero, but also a long tail (Simpson et al. 2017, Figure 2). The long tail accommodates the possibility of rapid changes in CPUE, such as a sudden decline in abundance that is caused by overfishing.

**Figure 2.**
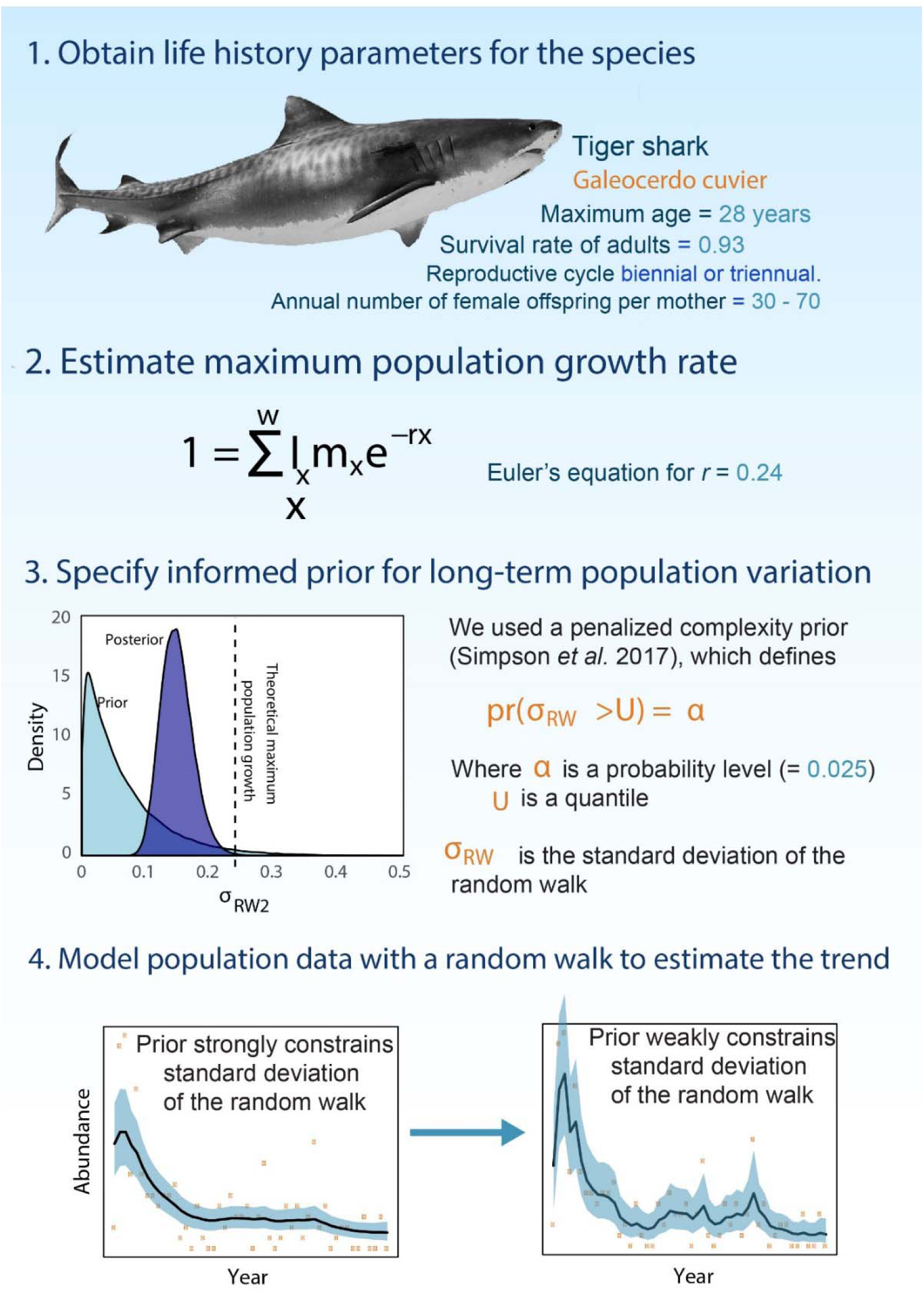
The approach for using life-history information to inform on trends in population modelling. The prior and posterior are shown for the fits to the full model with all data and the life-history prior.

Given that tiger sharks are large predators with relatively late age at maturity, a low fecundity and a high adult survival rate (Cortés 2002), we assumed that the instantaneous growth rate represents the near maximum rate of negative or positive annual change in the population. Therefore, we chose the penalized complexity prior such that there was only a 0.025 probability of a standard deviation greater than a prior estimate of *r* for tiger sharks. The exact probability level of 0.025 is arbitrary, but the probability should be small enough that annual changes larger than *r* are unlikely. We estimated *r* = 0.24 by applying the corrected Lotka-Euler equation (Cortés 2016; Pardo et al. 2016) to *G. cuvier* life-history parameters, (Supplementary Material), giving a prior probability of *r*_*t*_ > *r*_*max*_ (or *r*_*t*_ < −*r*_*max*_) = 0.016.

To explore whether informed priors could give more accurate results when sampling was limited to fewer regions, we compared results from the best-guess prior with three other priors. The first was the INLA default prior (a log-gamma with shape 1 and rate 1E-5)), which would represent a typical situation where the modeler does not use prior information on life-history. The second and third priors represented ‘bad guesses’ at an animal’s life history traits. They were penalized complexity priors parameterised to represent a very slow-growing shark species (*U* = *log*(1.02)) and a very fast growing shark species (*U = log*(1.66)) (Cortés 2002).

## Analyses

We first used simulation studies to address aim A, then analysis of the tiger shark data to address aims A-D.

### Simulation studies

We ran an initial analysis where we simulated random walks and then fit models to the simulated data. We simulated drifting random walks (with an average decline of 2.5% per year) that covered the range of shark species population growth rates with simulations for values of *exp*(*σ*_*r*_) = 1.02, 1.34 and 1.66 (Cortés 2002). For each trend and life-history type we simulated 20 replicate time-series of 30 years. Annual observations were then sampled from a negative binomial distribution with an expectation equal to the random walk value in each year and a dispersion (size) parameter = 2, to generate overdispersed counts. We repeated these analyses drawing observations from a Poisson distribution. We assumed effort was fixed over time and initial CPUE was set at 10 (approx. mean for tiger sharks). For each time-series we fitted three negative binomial models with different penalized complexity priors: U = ln(1.02), ln(1.34) and ln(1.66). In total we simulated 120 random time-series, fitting a total of 360 models. Each set of model fits was evaluated by: (1) the product of the likelihoods of the true (simulated) means in each year given the marginal probability distribution for the estimated annual means; (2) using the predictive ordinate as calculated by INLA, which is an in-sample measure of the model’s ability to predict the data (Held et al. 2010). For both evaluation measures lower values indicate the model does a poorer job at predicting the data.

We also studied the ability of different priors and models to detect sudden sustained population crashes. In these simulations we used the same priors and observation models as above and an initial CPUE of 50. We created time-series where the mean abundance was stable for 27 years, then collapsed in the 28th year to one of four values (12.5%, 25% or 50% of initial mean abundance). Models were fitted with random walks and either three or six years of post-decline observations (Supplementary Material).

### Analysis of tiger shark data

For the tiger shark data we first compare the complete model with two reduced models: one with an auto-regressive term on region only, which assumed that different gear types had similar temporal patterns, and one with no auto-regressive term and only a random intercept term for regional differences in mean catch. We fit all three model structures with both negative binomial and Poisson distributions. We used the predictive ordinate to choose between these models, visually verified model fits and then selected the best model for further analysis. We classified probability of decline according the IUCN Red List criteria A2, which for populations with unknown causes of decline have thresholds >80% (critically endangered), >50% (endangered), >30% (vulnerable) (IUCN 2017).

We then explore the impact of the extent of sampling and prior choice on estimates of decline over the time periods 1970 to 2017 and 1984 to 2017. We fit iterations of the best model to 60 subsamples of time-series drawn from different subsets of all regions and different prior specifications. We performed a factorial set of analyses crossing the four prior densities for *σ*_*RW*1_ with subsets of the number of regions included in the model fitting. We included 15 subsets of the set of 11 regions. The 15 subsets were a factorial cross of 1, 3, 6 and 9 regions crossed with a selection of regions grouped into: (A) the northern-most regions, (B) middle latitude regions, (C) southern-most regions and, (D) regions equally dispersed across the full extent of the dataset (fig. 1). For the subsets, we chose regions that had the most complete time-series. We also ran all priors for the complete set of 11 regions.

We compared results from all tiger shark model fits for their predictions of the magnitude of the population decline across two time-periods. Comparisons were made to the 11 region model with the life-history prior as our best-estimate. The % magnitude of population decline from the reference year was calculated as −100 (1 − *z*_2017_*/z*_0_) where *z*_2017_ was the value of the smoother in the most recent year and *z*_0_ was its value at the reference year (1970 or 1984). We used INLA’s ‘lincomb’ feature to calculate the marginal posterior distribution of the % decline statistic. We then compared scenarios for their median values and 95% credible intervals. We repeated the above analysis with a Generalized Additive Mixed Model (GAMM), so that the Bayesian method could be compared to more typical maximum likelihood methods (Supplementary Material).

## Results

### Simulation studies

In the simulation study, model fits were a more accurate representation of the true mean for slower growing species when compared to fast-growing species (Fig 3). For all species, the slow prior had a poorer fit to the data than the medium and fast priors, including, counterintuitively, for data simulated to represent a slow-growing species (Fig. 3). The slow prior was slightly poorer because it constrained the random walk too much to fully capture both year-to-year variation and the 2% per annum decline. The predictive ordinate, a standard in-sample evaluation measure did not detect any differences in the accuracy of fits by different priors (Fig S2).

**Figure 3.**
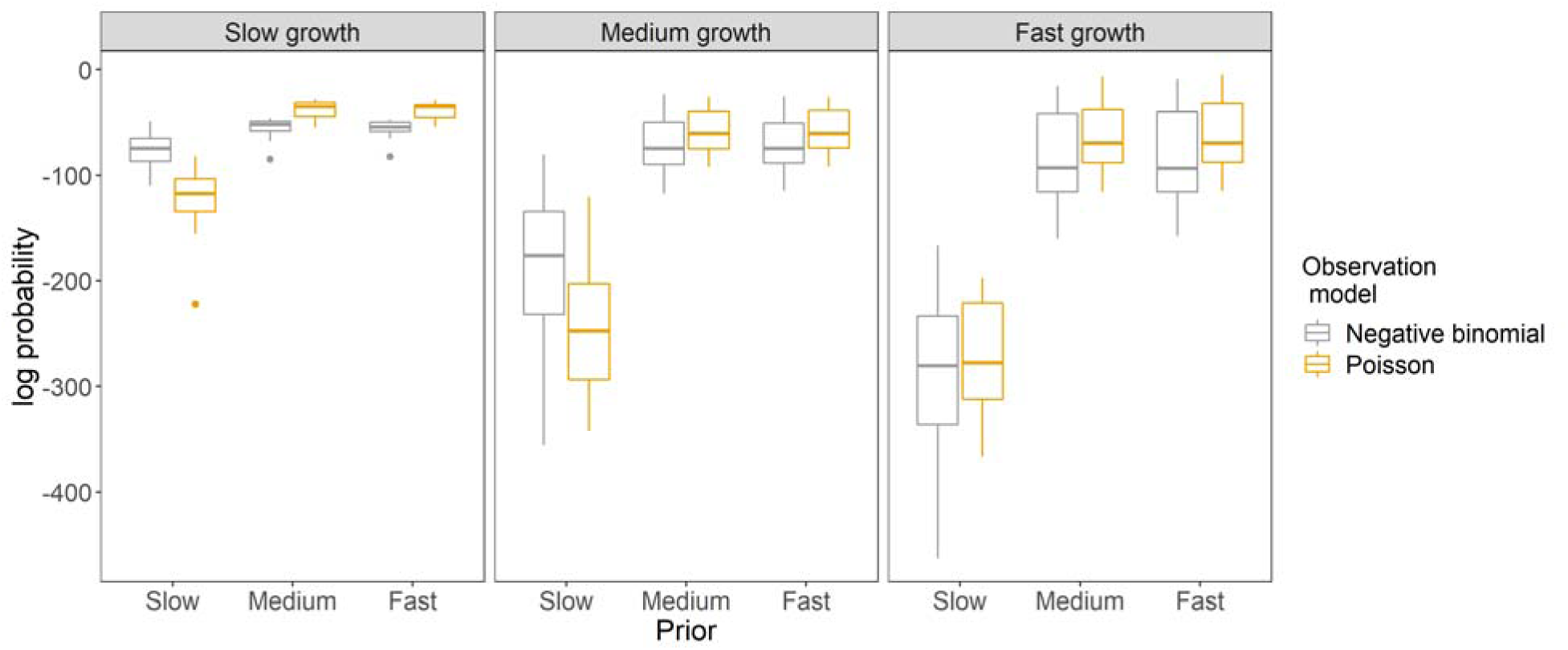
Results from simulation study for the log probability of the true simulated mean across all sample years given the results of the fitted model. Horizontal axes give the different priors, the panels show results from models fitted to time-series data simulated for species with different life-history traits. Higher (less negative values) indicate the model performed better at recovering the true trend. Boxes give the inter-quartile range and the horizontal bar gives the mean value. Vertical error bars extend no more than 1.5xIQR.

The simulation study of a sudden rapid decline showed that the faster priors were more likely to detect a sudden decline in CPUE (Fig S3). With the slow prior, the model estimated probability of a decline was always near 0.5. The models fitted to observations that were negatively binomially distributed were less sensitive to the decline than if the data were Poisson distributed. For instance, the probability of detecting a sudden decline with medium or fast priors was near one when data were Poisson distributed for any magnitude of decline and length of post-decline data. When data were negative binomially distributed the probability of decline was estimated to be between 0.55 and 0.88 for three years of post-decline data and between 0.6 and 0.9 for six years of post-decline data (Fig S3).

### Analysis of tiger shark data

For tiger sharks, the full model, with region by gear specific trends and a Poisson distribution had the highest predictive ordinate score (Table S2). Model verification measures also suggested this model best captured spatial and temporal variation in the data (Fig S4).

The estimated magnitude of decline for the east coast wide trend was greatest for the full model (71% over 3 generation), when compared to the simplified models (median estimates 40-61%). All models estimated a high probability that the population decline was greater than 50%, for the full model the probability was 0.96 (IUCN criteria A2 Endangered, Table S2). Interestingly, the full model detected regional variation in trends, with on average greater declines at the southern sites and less severe declines at the northern sites (Fig S6). We now proceed with further analysis using the full model.

For tiger sharks the magnitude of decline over three generations was similar with any prior and data from all regions (Fig 4, Fig 5). The slow prior tended to under-estimate non-linearities in the trend relative to the other priors and shrunk back to no trend when there were only data for a subset of regions (Fig 4D, Fig 5). The slow prior and the log-gamma priors tended to have the poorest coverage of the magnitude estimated with all when there were fewer regions included, they also had the highest risk of missed detection (Fig S7). The slow prior was also more confident about the magnitude of the trend (narrower credible intervals, Fig 5). Results were similar when estimating declines over 1970 - 2017 (Fig S8). The slow prior and the log-gamma priors gave results that were closer to those of the fast and life-history priors when the model did not include region specific trends in CPUE (Supplementary Material).

**Fig 4.**
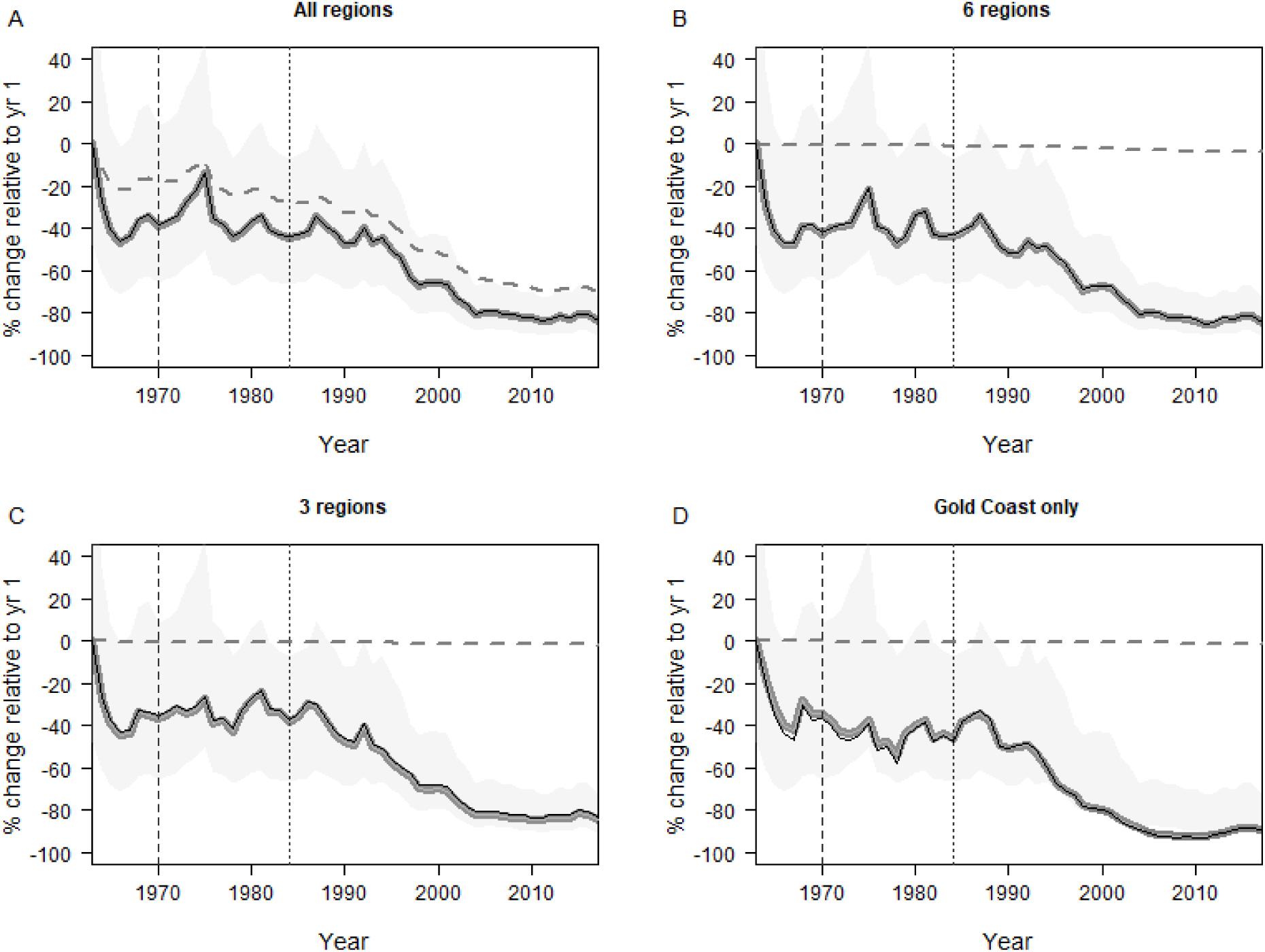
Examples of non-linear trends fitted to the tiger shark data. Shown is the east coast wide component of the trend from the full model fitted to all regions (A), and 6 and 3 extreme-latitude regions (B, C), and just the southern-most region (Gold Coast) (D). Lines show the fitted values for the random walk with 95% C.I.s (shading) for the life-history prior (thick grey line), slow life history prior (dashed) and INLA default prior (thin black line). Vertical dashed lines show baseline years used for analysis of % decline.

**Fig 5.**
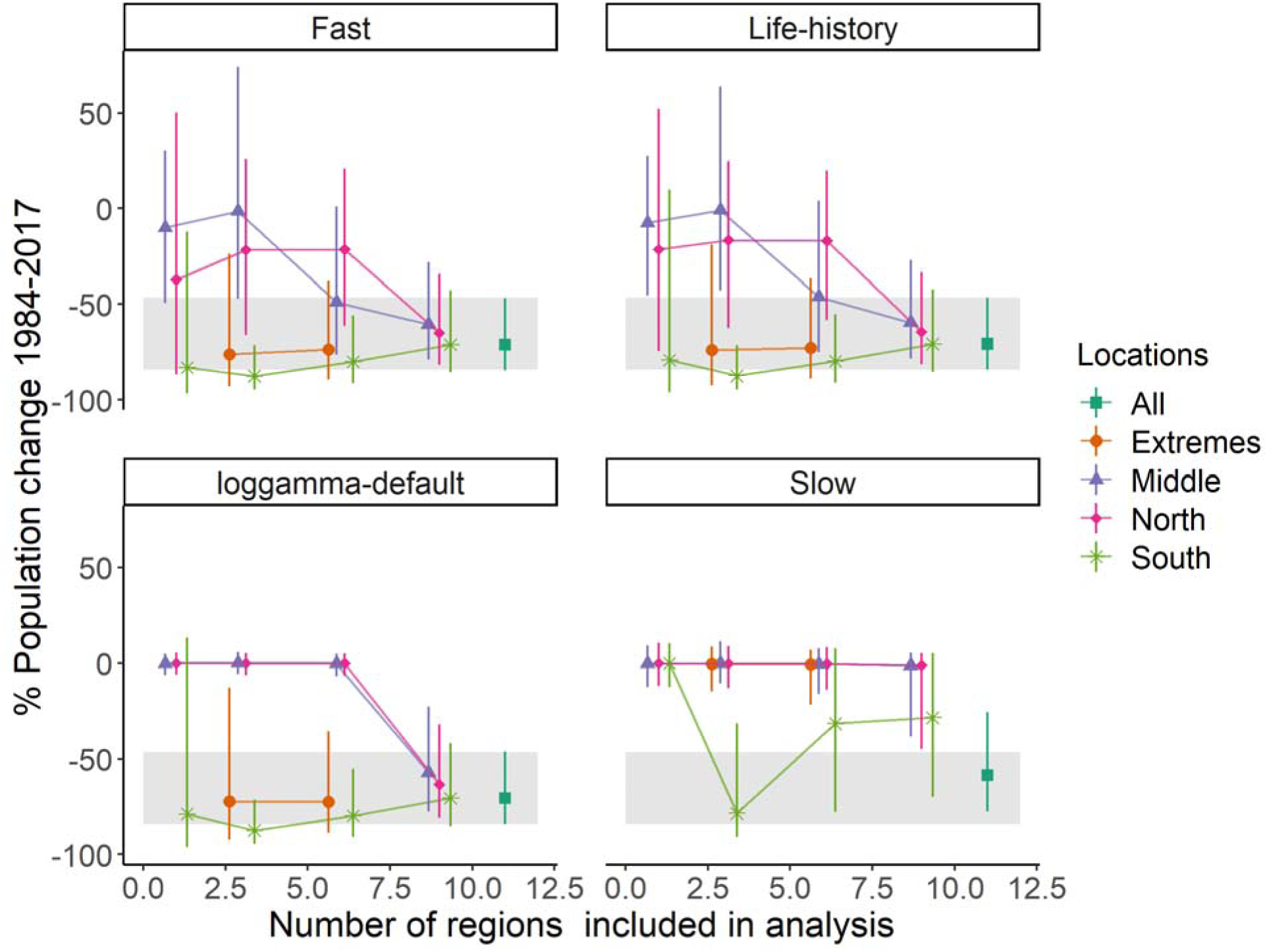
Estimated magnitudes of decline for the each prior (panels) and each scenario for subsets of regions (coloured points) over 3 generations (1984-2017). Points give median estimates and bars give 95% C.I.s. For comparison, the grey box shows the 95% C.I.s for the life-history prior fitted to data for all regions.

With a greater number of regions the estimate of decline converged to 71%. Subsets with southern regions had greater estimates of the decline, whereas subsets with middle and northern regions lower estimates of the rate of decline (Fig. 5). Data subsets that mixed extreme north and south regions were closer to the 71% decline than data subsets just of north or south regions. There was a notable spike in CPUE during the 1970s predicted by the model that used all regions and the life-history prior (Fig 4A), which was not present in subsets of the data (Fig 4B-D).

A generalized additive mixed model fitted using maximum likelihood methods showed a similar pattern to the Bayesian model of convergence of trend estimates as more regions were added, and greater declines estimated if data were taken from southern regions when compared to data taken from northern regions (Fig S9). The GAMM had similar results for either setting for its degrees of freedom.

## Discussion

### Predicting long-term population trends

We found that priors informed by life-history traits may improve the accuracy of population trend estimates. The use of life-history traits to inform on trend smoothing may help to overcome some of biases that come from analysing abundance indices with purely descriptive statistical models (Maunder et al. 2006). For instance, models that limit population growth within realistic bounds limit bias caused by observation errors (Sköld & Knape 2018). For the long-term tiger shark dataset, the estimate of decline was similar for all priors, so long as all sites were included in the analysis. The generalized additive model also estimated a similar rate of decline as the Bayesian model. The convergence in estimates across these different methods occurred because the tiger sharks’ CPUE data exhibited a strong trend. Where patterns in the data are strong, the prior will be less influential (e.g. Kindsvater et al. 2018). Despite the consistencies in trend estimates across the different methods, we still advocate using prior information to inform on population variability. Defensible parameter choices are important when model results may be contested, such as when governments make potentially contentious decisions about the status of populations (e.g. Edgar et al. 2018).

### Incorporating population declines into conservation

Conservation management must balance the risk of missing a true decline against the chance of a false alarm (Connors et al. 2014). In the tiger shark dataset, informed priors and sampling across more regions both improved the chances of detecting population decline. Calibration of the priors could be used to balance the trade-off between missed detection and false detection. For instance, our simulation study suggested that penalized complexity priors were more likely to detect sudden population collapses than priors that restricted variance. The shape of the penalized complexity prior was specifically designed to allow the data to speak for themselves when trends are strong, but to shrink estimates towards no trend when the data are weak or noisy (Simpson et al. 2017). A next step is to explore how calibrating the shape of the prior could be used to balance trade-offs between false alarms and missed detection.

Further research should test the method we propose across species with a broader range of life-history types. Life-history traits are useful predictors of trends across a broad range of species life-history strategies (e.g. Ribeiro et al. 2016), but classification errors for IUCN red list status tend to be greater for species with faster population growth (e.g. Sköld & Knape 2018). The approach proposed here will be most useful for detecting trends when temporary variation in abundance occurs over much shorter time-scales than variation in population size. For instance, the results of our simulation study were most accurate for slow-growing species that had less year-to-year variation, whereas accuracy was more variable for fast growing species that had greater year-to-year variation.

### Declines in tiger shark populations

The decline of tiger sharks we observed is consistent with a decline in large sharks throughout the world’s oceans (e.g. Baum & Myers 2004; Ferretti et al. 2008; Roff et al. 2018). The current global population trend for tiger sharks is unknown, although there is considerable variation across different oceans, with some regions showing no change (Baum & Myers 2004) and others large declines (Baum et al. 2003). The conservation status of tiger sharks globally escalated from “Lower Risk/near threatened” under the IUCN listing to “Near Threatened” in 2005 (Simpfendorfer 2009). Overall, their relatively high growth and reproductive rates (Cortés 2002; Holmes et al. 2015) means that tiger sharks are not considered at high risk of global extinction (Simpfendorfer 2009). However, eastern Australian and Oceania are a global hotspot of risk for tiger shark capture in long-line fisheries (Queiroz et al 2019). We found a high probability of a decline >50% on the east coast, which is consistent with listing tiger sharks as Endangered under state and national threatened species legislation. In nearby New South Wales tiger shark catches have also declined (Reid et al. 2011)), suggesting the decline of tiger sharks spans 18° in latitude across tropical and temperate coastlines of eastern Australia. This trend may also reflect a broader scale trend, as the east coast population is part of a well-mixed Indo-Pacific population (Holmes et al. 2017).

Across Queensland’s coastline we detected regional differences in the rate of decline, with more rapid declines in the Southern Queensland sites and slower declines or slight increases in the northern part of the range. Such rapid declines in southern regions are at odds with predictions of range extensions of tiger sharks from tropical to temperate coastlines of eastern Australian (Payne et al. 2018). However, the southward migration of tiger sharks in winter does expose them to heightened risk of capture in long-line fisheries (Queiroz et al 2019). Greater fishing pressure in more southern regions may interact with range edge dynamics to drive more rapid declines at the range extremes (Worm and Tittensor 2011). Further analysis of the tiger shark data could gain insights on the drivers of decline by studying how regional trends interact with the east coast wide trend. For instance, the magnitude of the regional auto-correlation terms represents the longevity of deviations from the broader trend. Thus, priors for the autocorrelation term could be modified to account for information on migration. Resolving the drivers of trends in tiger sharks is important to inform on conservation actions.

## Conclusions

In our models, regional differences in population trends meant the accuracy of estimates of decline was improved when data from a greater number of regions was available. However, spatially extensive sampling can be expensive and not all species will have spatially extensive time-series available for assessing trends (Kindsvater et al. 2018). We recommend that where possible estimates of population change for wide-ranging species should use data from a large part of the species’ range. We considered how estimates of population trends for wide-ranging species depend both on the spatial extent of data and the model used to partition short-term variation from long-term trends in population size. We found that the choice of model and the spatial extent of sampling interact to effect estimates of population trends. Our results provide further support for the use of life-history information to help overcome the deficiency of monitoring data that exists for many potentially threatened species (Kindsvater et al. 2018). We suggest this approach is complementary to existing population modelling approaches and may be particularly useful for rapid assessments of population trends.

