## Supplementary Material for "Life-history traits inform on population trends when assessing the conservation status of a declining tiger shark population"

### Estimating the instantaneous rate of increase for *G. cuvier*

We estimated an instantaneous rate of population increase for Pacific *G. cuvier* of 0.24. This was based on applying the Euler-Lotka equation to life-history parameters and is consistent with previous estimates for tiger sharks from the Atlantic ocean (e.g. Cortés 2016, Pardo et al. 2016)). Our estimate was based on the closest available life-history parameters: an age at maturity of females of 12 years (east coast of Australia, Holmes et al. 2015), an average female fecundity of 32.6 pups every 3 years (Hawai’i, Whitney & Crow 2007) a lifespan of 28 years (Cortés 2016) and an instantaneous survival rate of 0.93 per year (Cortés 2016). Differences in life-history parameters across regions will not affect our conclusions because the large amount of data meant the results were not sensitive to small changes in r and thus prior specification.

### Prior choice

We specified prior distributions for the parameters $\alpha$, $\beta$ and the hyper-parameters $\theta$, $\rho$ and $\sigma_{AC1}$. For $\alpha$ and $\beta$ we used broad normal priors with N(0, 0.0) and N(0, 0.001), respectively (the defaults for the software we used, Martins et al. 2013). For $\theta$ we used the penalized complexity prior with one parameter, set to 7 based on a simulation study (Supplementary Material). The penalized complexity prior will shrink the negative binomial distribution toward a Poisson distribution if there is not strong evidence for over-dispersion ($\sigma^{2}>>\mu$) (Simpson et al. 2017). For the region specific autoregressive terms we also used the default penalized complexity priors for $\rho$and $\sigma_{AC1}$, which weakly shrink estimates towards $\rho=0$ and $\sigma_{AC1}$=0 (and Sørbye and Rue 2016). Using weakly informative priors means computations are more efficient and avoids overfitting the data to the variance parameters (Simpson et al. 2017).

The second reduced model for tiger sharks had only a time-constant random effect by region (called the ‘iid’ or independent random noise model in the INLA documentation). For this variance parameter we used a weakly informative log-gamma prior on the precision (=1/variance) with parameters shape = 1 and rate = 1e-5.

### Further details of INLA settings

In the INLA implementation we set the ‘scale.model’ option equals ‘FALSE’ to ensure the value provided for $\sigma_{RW1}$ was interpreted as the marginal standard deviation. We also set ‘constr’ equals ‘TRUE’, although this choice has no effect on our results or interpretation. Setting ‘constr = TRUE’ means we could estimate an overall model intercept, which represents average catch across all regions and years. Setting ‘contstr = FALSE’ has no effect on model predictions, but would just mean we could not separate an intercept term from the random effects of regions and years.

We estimated per cent declines using INLA’s ‘lincomb’ feature. Further details of model implementation are available on the github site.

### Determination of priors for the size parameter of the negative binomial

For the prior for the negative binomial dispersion parameter we used the penalized complexity prior for $\theta$ as implemented in the INLA program. Because this prior is relatively new, we ran a simulation study to choose its parameterisation. Note that INLA’s parameterization of the negative binomial has the variance grow exponentially with its mean.

For the simulation study we randomly generated 30 random walks of 40 years each, under two assumptions. In the first, observation errors were sampled from a Poison distribution (i.e. $\theta=\infty$. In the second, observation errors were sampled from a negative binomial distribution with $\theta=2.1$, which approximates the value estimated in the model with all regions and indicates a moderate level of over-dispersion. We then fit the models to each time-series with three prior values of 1, 7 (the default) and 20 for the penalized complexity gamma prior ‘pc.mgamma’ (Martins et al. 2013). Higher parameter values for the prior place greater prior density on overdispersion (low $\theta$). We then evaluated the model fits to the two types of distributions by estimating the proportional coverage of the 95% CIs over the true dispersion value, where an unbiased model should have coverage of approximately 95%.

The model with $\lambda=7$ had the highest coverage for the negative binomial data (Table S1, Fig S1). All Poisson models have reasonable coverage of values >30 (which will be approximately Poisson distributed).

**Table S1** Coverage of the ‘true’ parameter value by each model.

| distribution | lambda | coverage |
| --- | --- | --- |
| negbin | 0.1 | 0.97 |
| negbin | 7.0 | 0.97 |
| negbin | 20.0 | 0.90 |
| pois | 0.1 | 0.83 |
| pois | 7.0 | 0.93 |
| pois | 20.0 | 0.97 |

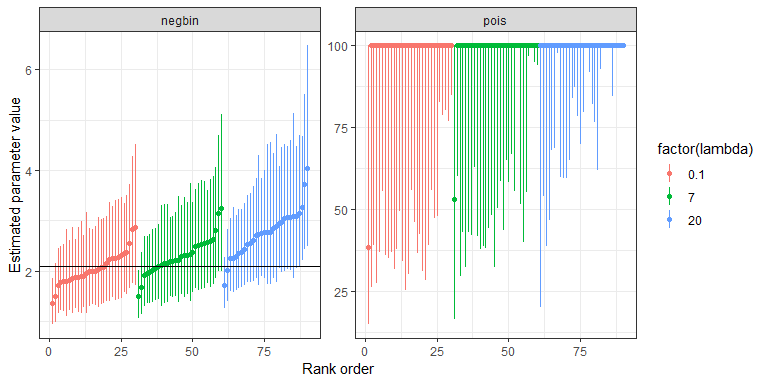

**Fig S1** Sensitivity analysis for estimates of the negative binomial overdispersion parameter when fitting a random walk model to negative binomially distributed data (A) or Poisson distributed data (B). Coloured points and errorbars show individual means and 95% C.I.s for randomized data-sets fit with different priors. The solid black line in (A) shows the ‘true’ value of the dispersion parameter for the negative binomial. For the Poisson, larger values indicate the model estimated a posterior closer to a Poisson distribution.

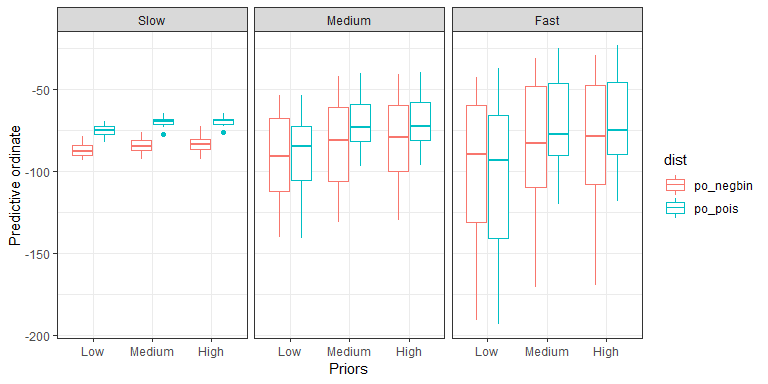

**Fig S2** Results for the predictive ordinate for the simulation study. Horizontal axis shows different priors, the panels show results from models fitted to data simulated for different species traits and the colours show data simulated from Poisson and negative binomial distributions.

### Testing the ability of models to detect sudden rapid declines

The life-history priors imply symmetrical probabilities of population increase and decreases, so may influence our ability to detect sudden rapid declines. We created a time-series where the mean abundance was stable at 50 individuals for 27 years. In the 28th year we forced a sudden decline to 12.5%, 25% or 50% of the initial value. We then created two post-decline periods of three years and six years. We simulated twenty observed time-series each with for Poisson and negative binomial errors (theta = 2). To each simulated series we fitted negative binomial Bayesian models with a random walk order one on the temporal effect. We fitted the model three times to each time-series with each of the three priors used in the main text. In total we fitted 720 models (three priors by two types of observation error by twenty replicates by two lengths of post-decline periods by three levels of decline).

We evaluated the outcomes of the simulations by estimating the decline from the 27th year to the final year of each time-series with 95% credible intervals. For each simulation and prior setting we then calculated the average probability of zero decline or greater.

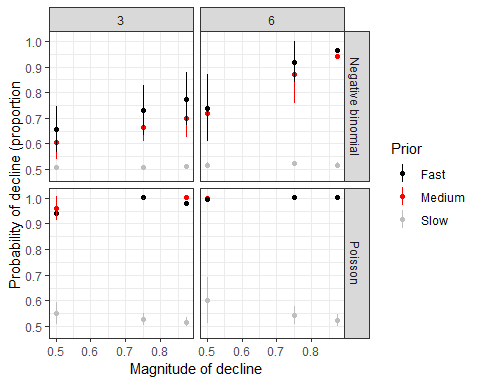

**Fig S3** Model estimated probabilities that abundance declined in the 27th year of simulated time-series. Random walk models were fitted to replicate time-series with different true declines (x-axis), three priors (colours), Poisson or negative binomially distributed data (rows) and three or six years post-decline data (columns).

### Model comparison for tiger shark data

**Table S2** Results from the six models fitted to the tiger shark data, all models have a fixed gear effect and an east coast wide trend estimated with a random walk (equation 2). 95% credible intervals are given in brackets. Lower value of the WAIC indicate a more parsimonious model, higher (less negative) values of the predictive ordinate indicate greater predictive ability of the model.

| **Model** | **Distribution** | **Magnitude of decline over 3 generations** | **Probability of decline >30%** | **Probability of decline >50%** | **Probability of decline >80%** | **WAIC** | **Predictive ordinate** |
| --- | --- | --- | --- | --- | --- | --- | --- |
| Random intercepts for regions | Poisson | 50% (42-59) | 1 | 0.57 | 0 | 8344 | -3900 |
| Random intercepts for regions | Negative binomial | 61% (46-71) | 1 | 0.93 | 0 | 6023 | -2984 |
| Region specific temporal auto-correlation | Poisson | 57% (24-75) | 0.95 | 0.69 | 0 | 7308 | -2939 |
| Region specific temporal auto-correlation | Negative binomial | 58% (33-74) | 0.98 | 0.77 | 0 | 5879 | -2866 |
| Gear by region specific temporal auto-correlation | Poisson | 71% (46-84) | 1 | 0.96 | 0.11 | 4832 | -2066 |
| Gear by region specific temporal auto-correlation | Negative binomial | 71% (48-84) | 1 | 0.97 | 0.1 | 5074 | -2258 |

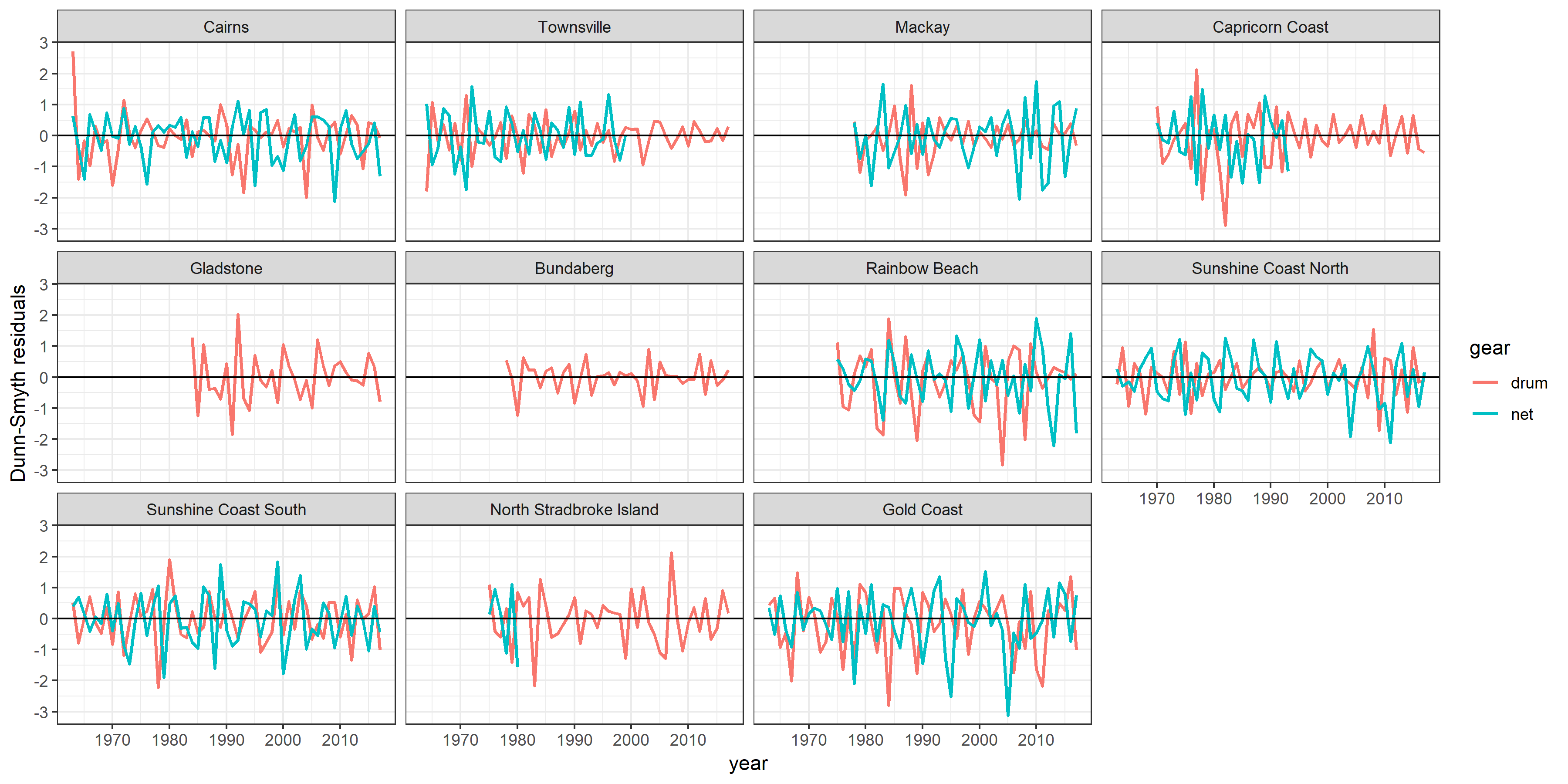

**Fig S4** Dunn-Smyth residuals (Dunn and Smyth 1996) by year, region and gear type for the complete model.

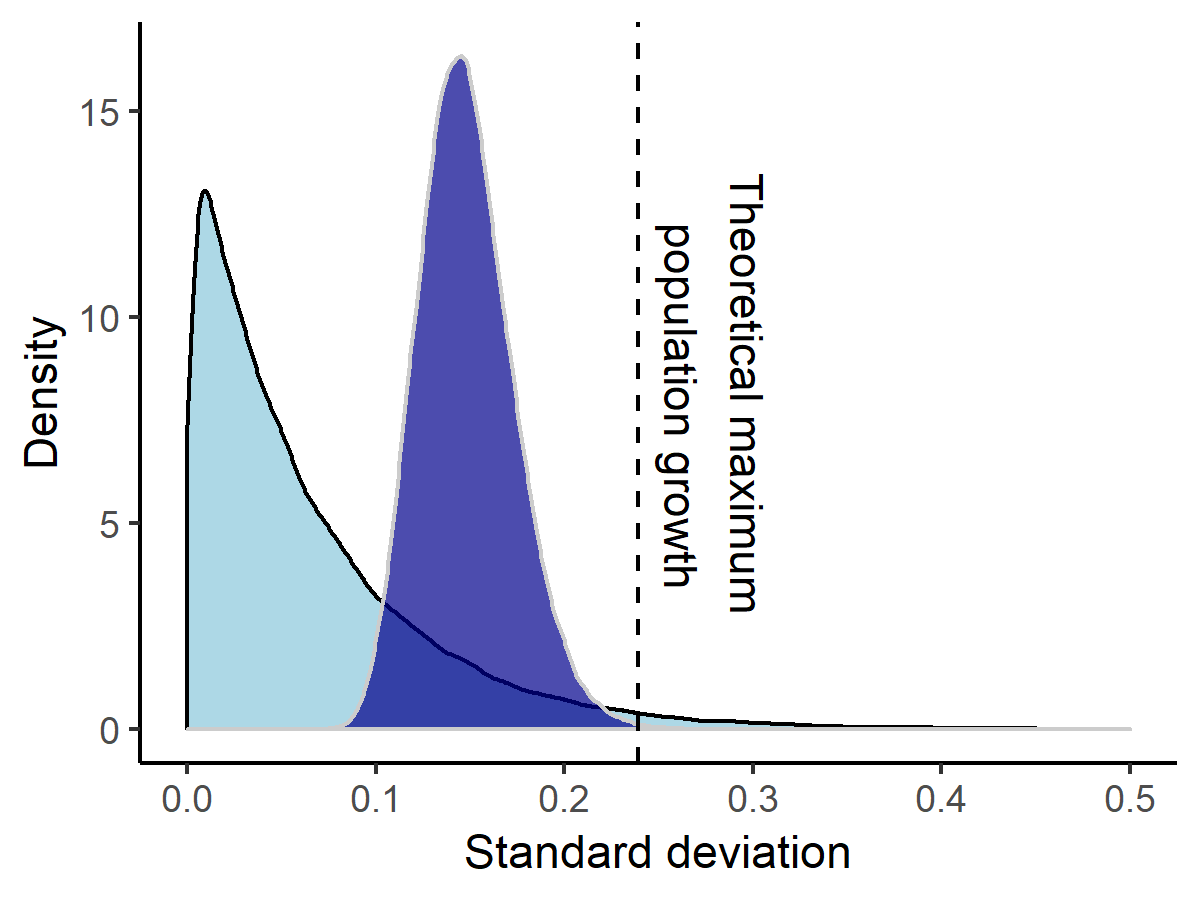

**Fig S5** Prior (light shading) and posterior (dark shading) estimates for the standard deviation of the random walk. Threshold for the maximum population growth rate with a <0.025 probability is shown with dashed line.

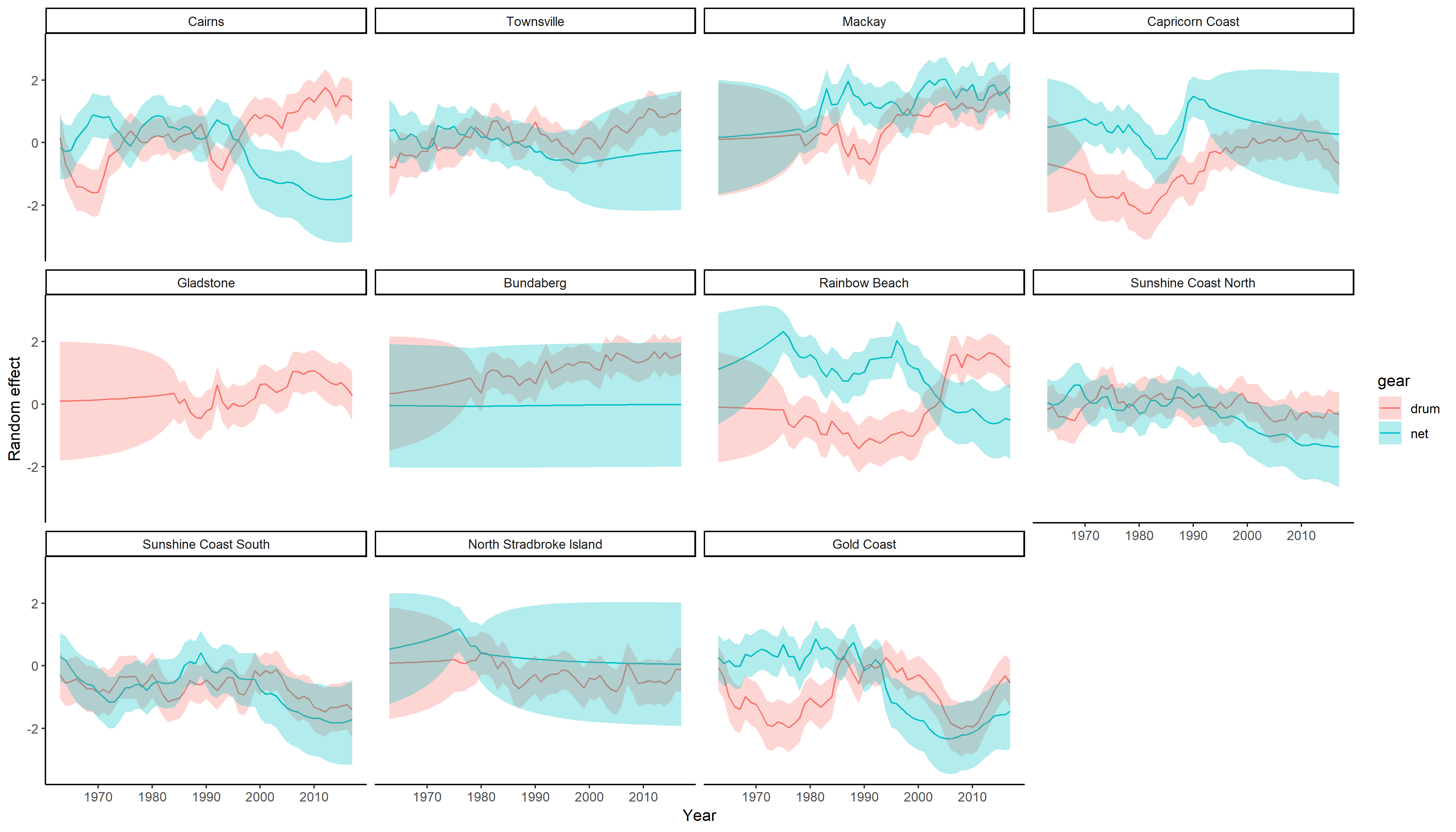

**Fig S6** Random effect estimates for the region by gear autoregressive term. Shown are the median (solid line) with 95% credible intervals (shading) on the log-link scale. The random effects estimates show deviations from the east-coast wide trend (random walk) and are analogous to partial effects (so the total effect would be the sum of the autoregressive and random walk components). Regions are ordered north to south from top-left to bottom-right.

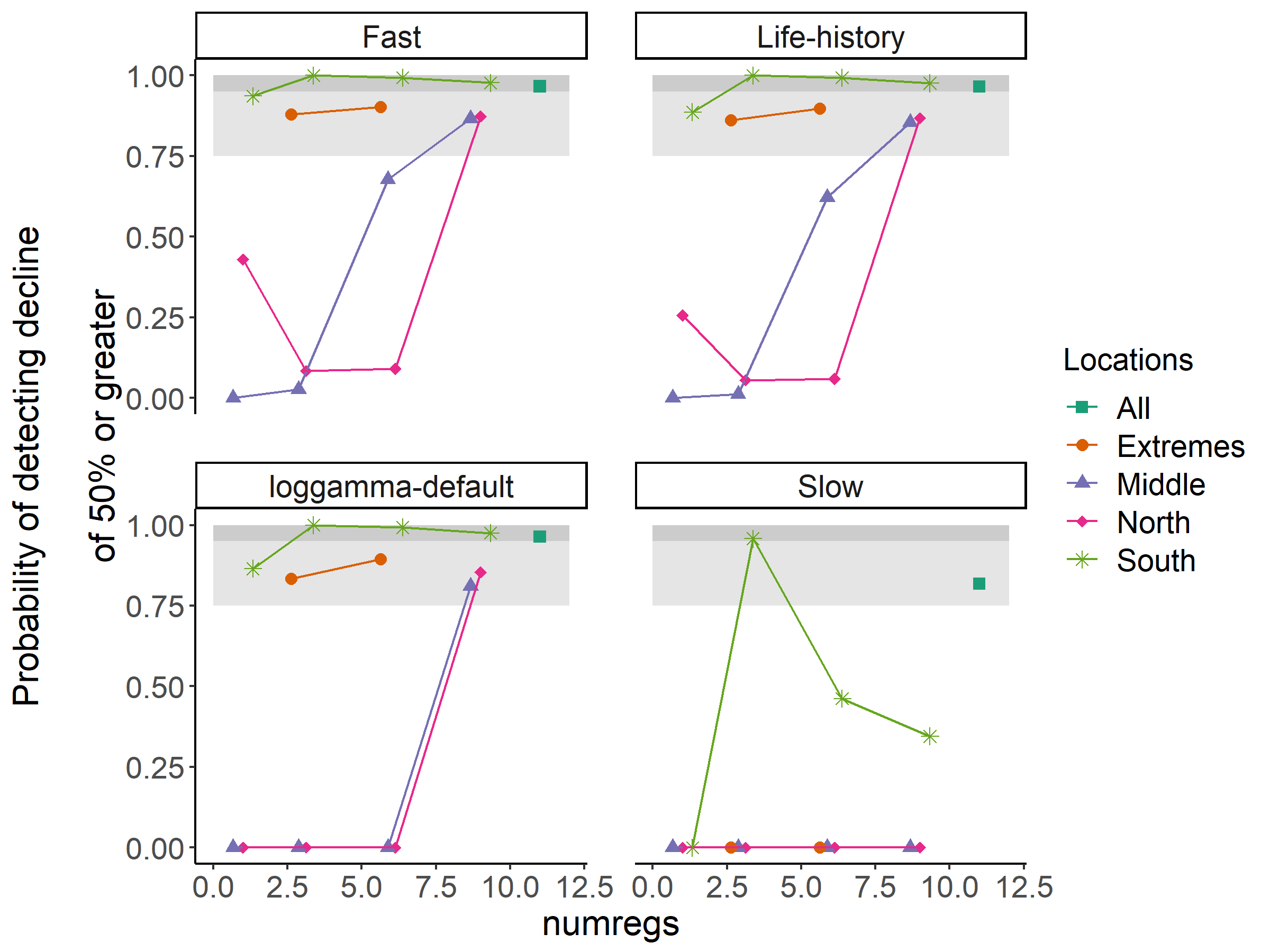

**Fig S7** Model power measured as probability of detecting a decline of 50% or greater, for the each prior (panels) and each scenario for subsets of regions (coloured points) over 3 generations (1984-2017).

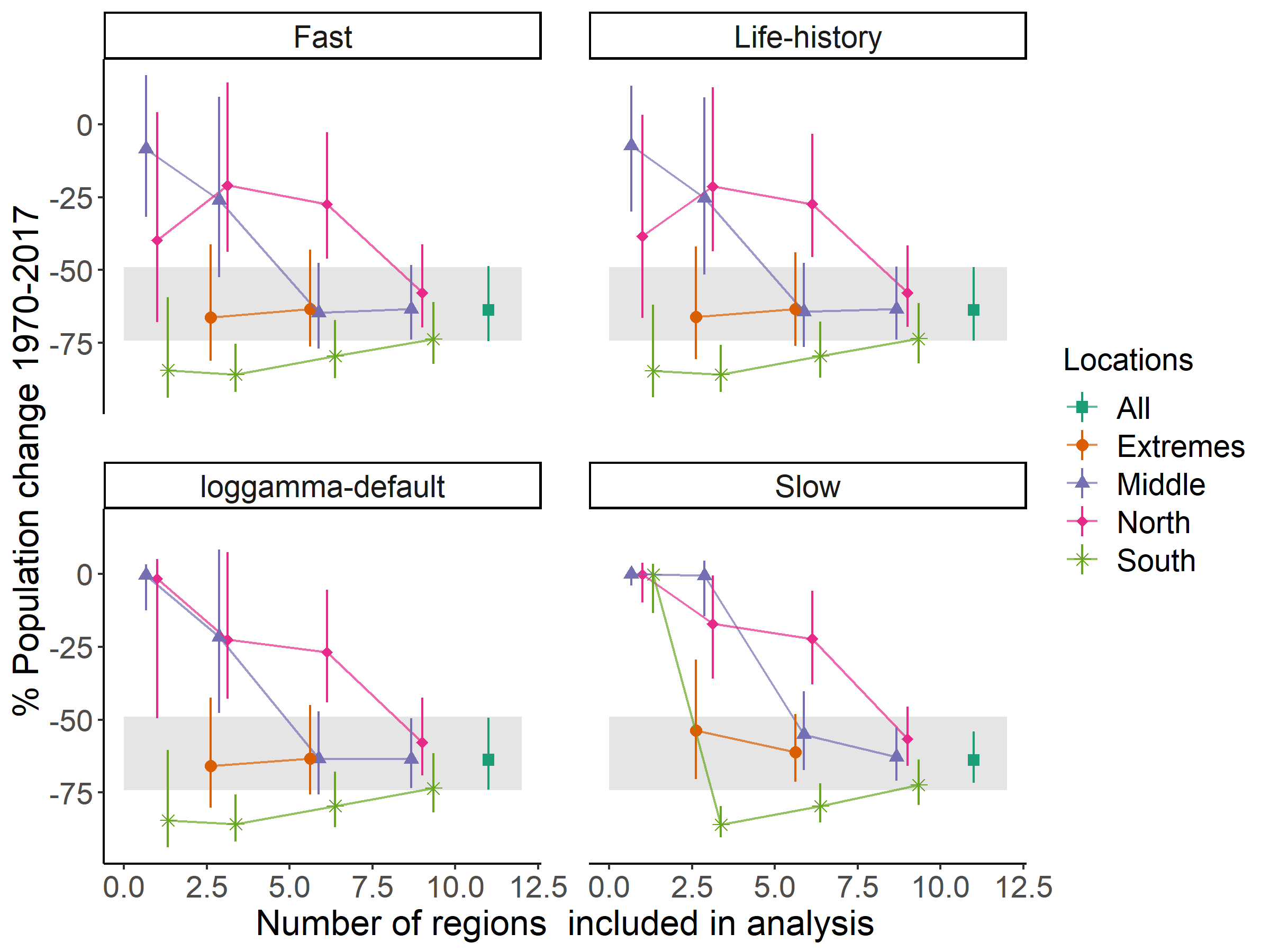

**Fig S8** Estimated magnitudes of decline for the each prior (panels) and each scenario for subsets of regions (coloured points) over 1970-2017. Points give median estimates and bars give 95% C.I.s. For comparison, grey region shows the 95% C.I.s for the Bayesian model with the life-history prior and fitted to data for all regions.

### Comparison of Bayesian models to Generalized Additive Mixed Models

We compared the main results to those generated by a Generalized Additive Mixed Model (GAMM, fitted with the R package mgcv, version 1.8-23 Wood 2017) with a negative binomial distribution for the data. The negative binomial parameter theta parameter was fixed at a value of 2.5, based on exploratory fitting (Wood 2017). The GAMM was specified to be equivalent to the full INLA model. It had random effects by regions (for sub-samples with >1 region), a thin plate smoothing spline applied to year, a fixed effect for gear type and an offset on log effort. We included an auto-regressive order 1 (AR1) correlation structure, where the AR1 term was applied by regions and gears. The models fit with only one region excluded the AR1 effect. The maximum degrees of freedom was set to either 1/3 the number of years (Fewster et al. 2000), or chosen with cross-validation (Wood 2017). The model code for the GAMM was thus given as:

gamm(abundance ~ s(yearnorm, k = k_upper)+ gear + offset(Leffort), correlation = corAR1(form =~ yearnorm2 | regfact), family = negbin(theta = theta_fixed))

Where ‘regfact’ is a factor of gear by region combinations and ‘theta_fixed = 2.5’.

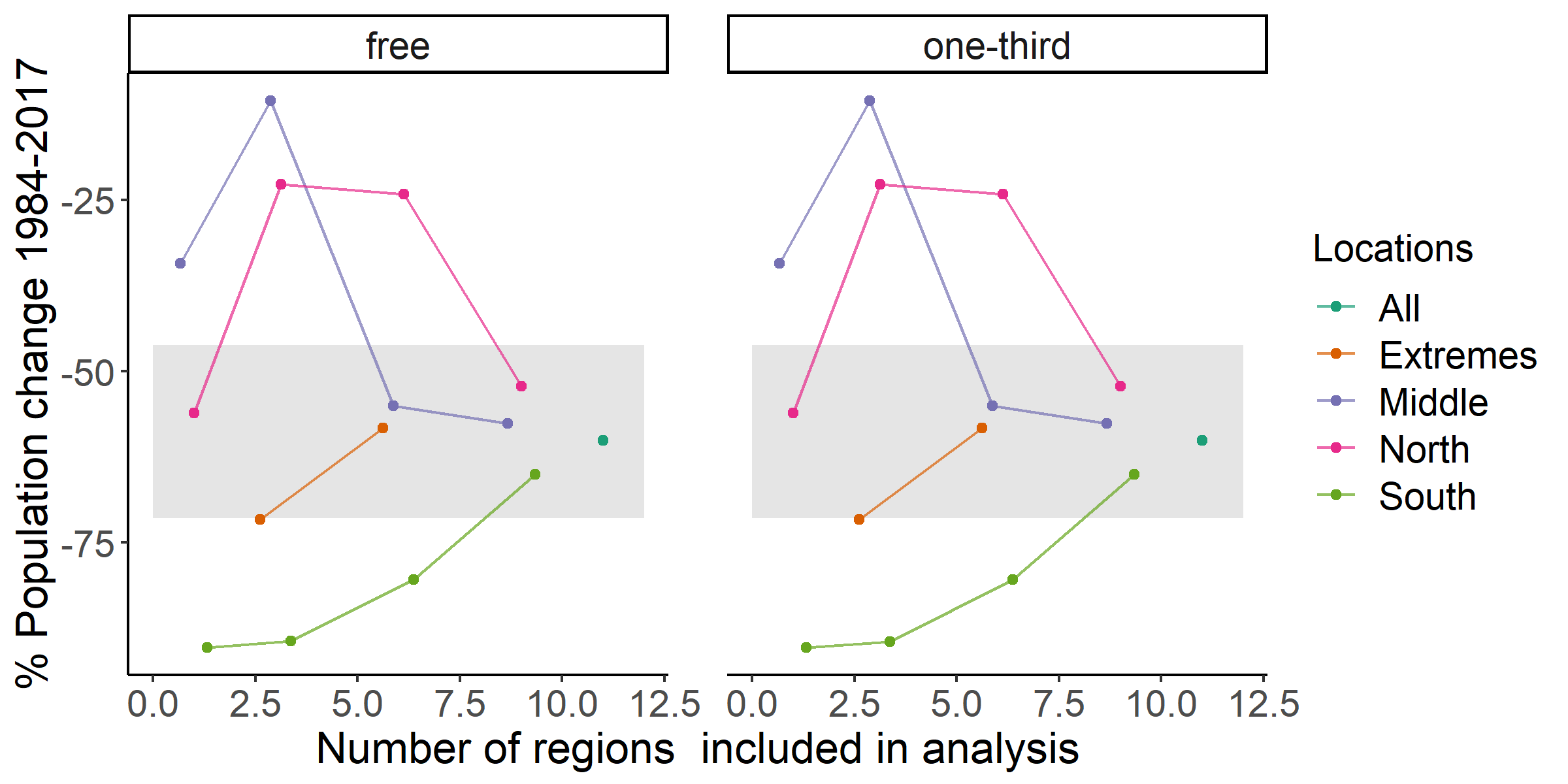

**Fig S9** Estimated magnitudes of decline from a generalized additive mixed model with (A) cross-validated estimation of degrees of freedom and (B) maximum degrees of freedom set to 1/3 the length of the time-series. Horizontal grey lines show estimated decline from the Bayesian model.
